# Sex difference in the facilitation of fear learning by prior fear conditioning

**DOI:** 10.1101/2023.06.29.547102

**Authors:** Kehinde E. Cole, Ryan G. Parsons

**Affiliations:** Stony Brook University, Department of Psychology, 100 Nicolls Rd., Stony Brook, NY, 11794

**Keywords:** Learning, Memory, Fear, Sex difference, Metaplasticity, Tagging, Allocation

## Abstract

There is now ample evidence that the strength and underlying mechanisms of memory formation can be drastically altered by prior experience. However, the prior work using rodent models on this topic has used only males as subjects, and as a result, we do know whether or not the effects of prior experience on subsequent learning are similar in both sexes. As a first step towards addressing this shortcoming rats of both sexes were given auditory fear conditioning, or fear conditioning with unsignaled shocks, followed an hour or a day later by a single pairing of light and shock. Fear memory for each experience was assessed by measuring freezing behavior to the auditory cue and fear-potentiated startle to the light. Results showed that males trained with auditory fear conditioning showed facilitated learning to the subsequent visual fear conditioning session when the two training sessions were separated by one hour or one day. Females showed evidence of facilitation in rats given auditory conditioning when they were spaced by an hour, but not when they were spaced by one day. Contextual fear conditioning did not support the facilitation of subsequent learning under any conditions. These results indicate that the mechanism by which prior fear conditioning facilitates subsequent learning differs between sexes, and they set the stage for mechanistic studies to understand the neurobiological basis of this sex difference.

## Introduction

When determining how the brain acquires new aversive memories, laboratory studies have most often used learning experiences that are isolated from other experimentally-controlled events. This approach has been valuable, but learning does not typically occur in isolation from other experiences, and in fact, many recent studies have described several conditions in which prior experience can alter the strength and underlying mechanisms of learning. For example, our previous work using Pavlovian fear conditioning showed that a single pairing of light and mild shock, insufficient to support memory formation when presented alone, was able to prime learning when a second trial was later presented (Parsons and Davis, 2012; Parsons, Walker and Davis, 2016; Cole et al., 2019). This priming effect was time dependent such that optimal facilitation was observed when the second trial occurred within an hour to several days later. More recently, we showed that prior auditory fear conditioning was similarly able to facilitate long-term memory formation when rats were given a single pairing of light and shock an hour or 1 day after auditory fear conditioning (Lee et al., 2018). Analogous facilitation-like effects between separate fear conditioning events have been reported by others as well (e.g., Rashid et al., 2016), and more broadly, there is a growing literature describing conditions in which prior learning can alter subsequent memory formation.

While there is now good evidence that prior fear conditioning can facilitate future fear learning, there is no prior work testing whether there are differences in males and females. In addition, while there are numerous reports of facilitation effects between two distinct cued fear conditioning events, it is unknown whether or not contextual fear conditioning primes future memory in the same manner. Answering these questions is important from the broad perspective of how sex affects the basic mechanisms of learning and memory, and it is also important with respect to disorders of fear and anxiety, which often carry a higher incidence in human females (Kessler et al., 1995; Kilpatrick et al., 2013), and are characterized in part by dysregulated fear learning (Parsons and Ressler, 2013).

In our prior work in males, we showed that a typical auditory fear conditioning session was able to facilitate future visual fear conditioning if the subsequent training occurred 1 hour or 1 day later (Lee et al., 2018). In the work described here, we tested whether or not males and females would differ in this paradigm if the two training sessions were separated by an hour or 1 day. In addition, we also tested whether or not unsignaled shocks (i.e., contextual fear conditioning) would facilitate subsequent visual fear conditioning. Fear behavior to both cues was examined in separate test sessions following the two training events.

## Materials and Methods

All procedures were approved by the Stony Brook University Institutional Animal Care and Use Committee and were in accordance with the National Institutes of Health guidelines for the care and use of laboratory animals.

### Subjects

A total of two-hundred and fifteen, male (n = 107, 300g – 325g upon arrival) and female Sprague-Dawley rats (n = 108, 200g – 225g upon arrival), at least 8 weeks of age obtained from Charles River Laboratories (Raleigh, NC, USA) served as subjects. Rats were housed in pairs in a colony room with a 12h light-dark cycle with lights on at 0700 h. Rats had food and water provided freely throughout the experiment. On delivery, rats were left undisturbed for 7 days, and then each rat was gently handled for 5 min every day for 6 days. During the last 3 days of handling, rats were carted into the laboratory and handled to acclimate them to being transported. All behavioral procedures began after the sixth day of handling.

### Behavioral Apparatus

#### Baseline acoustic startle response and Fear-Potentiated Startle

Startle amplitude was assessed using a Startle Monitor II system (Kinder Scientific, Poway, CA; Version 8.15). Baseline startle sessions and test sessions occurred in a set of four identical 17.5 × 9.2 × 7.5 cm restrainers which sat atop a load cell sensor within a (40.64 x 40 x 49.53 cm) sound attenuating chamber. Visual fear conditioning occurred in a set of four identical 26.67 x 20.96 x 15.9 cm (depth x width x height) plexiglass and stainless-steel cages. The floor of these cages was made of stainless-steel bars through which shock could be delivered. Each of these steel cages sat atop load sensors and housed individually within sound attenuating chambers. Startle responses or movement caused movement of the restrainer detected by the load cell and transduced into a voltage change that was then converted to Newton (N) through a single-pulse calibrator interfaced to a PC. Speakers were located on the ceiling of each chamber through which a white noise burst was delivered (50 ms, 95 dB) to elicit the startle response. The same speakers produced a constant background noise of ∼54 dB. The light (4.0 s/82 lux) stimulus was delivered through an LED light panel positioned on the ceiling of the cabinets. Startle amplitude was defined as the peak N that occurred during the 500 ms following the onset of a white noise burst. Cages were wiped down with 70% EtOH to provide an olfactory cue.

#### Freezing

Rats underwent auditory fear conditioning in a set of four identical 32 x 26 x 21 cm conditioning chambers made of stainless steel and Plexiglass walls, a shock grid floor located within a sound-attenuating 45.7 x 43.2 x 43.2 cm isolation boxes, (Clever Sys. Inc., Reston, VA; CSI-BOX-STD). During training (Context A) 28-V, incandescent, house light bulbs were used, and the walls of the context were wiped down with 5% acetic acid to provide an olfactory cue. Overhead cameras recorded behavioral sessions and the video signal from each chamber fed into a software program (FreezeScan 2.0) which automatically scored freezing behavior based on pixel change. On testing day, the context (Context B) was altered by adding painted inserts to the floor and walls of the chambers. A bent 33.5 x 21.3 cm metal insert was placed into each of the chambers, changing their shape. Before testing, the chambers were wiped down with 5% ammonium hydroxide to provide a different olfactory cue from training day. In addition, rats were transported in buckets from the colony room into the lab instead of their home cages, and LED light was used to illuminate the chambers instead of the house light bulbs. Freezing behavior was measured as an index of fear learning and was defined as making no movement other than what was necessary to breathe.

### Behavioral Procedures

#### Baseline Startle

All subjects underwent baseline startle amplitude procedures on two consecutive days. Rats were placed in startle cages, and after a 5 min habituation period, rats were presented with 30, 95 dB, 50 ms, white noise bursts (30 s ITI). Startle amplitude was defined as either the peak N that occurred during the 500 ms following the onset of a white noise burst or by displacement of an accelerometer that produced a voltage output proportional to the velocity of cage movement during the first 200 ms after onset of the startle-eliciting noise bursts. The mean startle amplitude across the 2 days were determined for each rat, and animals were assigned to groups (Tone-shock, Shock-alone, or No light-shock) that had equivalent startle amplitudes.

#### Auditory/contextual fear conditioning

The following day after baseline startle, rats were exposed to an auditory or contextual fear conditioning procedure. Rats in the Tone-shock (n = 18 females, n = 18 males) and No light-shock (n = 18 females, n = 18 males) groups were placed into the freezing chambers (Context A) and given two pairings of 30 s, 4kHz 76dB tone (CS), co-terminating with a 1 s, 1 mA footshock (US) (2 min ITI) following a 6 min baseline habituation period. Rats in the Shock alone (n = 18 females, n = 18 males) group received two 1s, 1 mA footshocks (2 min ITI) in the absence of a tone following the 6 min baseline habituation period. All rats remained in the freezing chamber for an additional two minutes after the final shock and were then returned to their home cages.

One or twenty-four hours after the auditory fear conditioning procedure, rats underwent visual fear conditioning. Rats were placed in shock cages in the acoustic startle chambers and presented with a single light (4 s, 82 lux)-shock (0.5 s/0.4 mA) pairing following a 5 min baseline habituation period. After the light-shock trial, rats remained in the chambers for an additional two minutes after the shock before they were returned to their home cages. All rats that had been exposed to contextual fear conditioning (n=38 males, n=36 females) the prior day received the light-shock training. One-half of the animals (n=34 males, n = 36 females) that had received auditory fear conditioning were also trained in the light fear conditioning procedure, while the other half (n=36 males, n=36 females) were placed in the shock cages for 7 min with no presentation of the light or shock.

#### Fear memory Testing

Rats were tested 48 h after the final training session. Rats were tested first for fear to the visual cue and then tested to the auditory cue the next day. Rats were tested to the light cue in the chambers used during the baseline acoustic startle sessions. After a 5 min baseline period, rats were presented with 30 startle trials to habituate startle responses prior to the onset of the light-startle trials. Rats then received 40 additional trials, consisting of 10 light-startle trials, each followed by 3 startle-alone trials. For the light-startle trials, the 95 dB white noise burst was presented 3.5 s after onset of the cue. For the startle-alone trials, the 95 dB white noise burst was presented in the absence of the light. Rats were returned to their home cages 2 mins after presentation of the last startle-alone trial.

Twenty-four hours after testing to the visual cue, rats were then tested in the freezing chambers in Context B. Following a 6 min acclimation period, rats were presented with 8 trials of the auditory cue in the absence of the shock, with a 2 min inter-trial interval between each presentation. Rats were returned to their home cages 2 mins after presentation of the last tone.

### Data analysis

Baseline startle amplitude for each rat was calculated by taking the average of the startle responses across the 2 days of baseline startle testing. These values were used to match rats into groups with equivalent mean startle amplitude within each sex. Fear behavior during the tone fear conditioning and testing sessions was calculated by taking an average of the total time spent freezing during the 6 min acclimation period, and the average time spent freezing during the CS periods. Shock reactivity during the light fear conditioning session was measured as the peak change in force that occurred during the shock period. Fear-potentiated startle during the test sessions was calculated by finding the average of the 10 light-startle trials and of the 30 startle-alone trials for each rat and calculating a percent change score. Percent change scores were used (i.e., vs. absolute difference scores) because of previous work showing that they remain stable across large variations in baseline startle amplitude (Walker and Davis 2002). Baseline fear during light testing was assessed by subtracting average baseline startle amplitude for each rat from the average amplitude of the 30 startle trials at the beginning of testing and calculating a percent change score. Likewise, fear during the intertrial interval was computed by subtracting the average amplitude of the 30 startle trials at the beginning of testing from the average baseline startle amplitude for each rat and calculating a percent change score.

Statistical analyses were performed using SPSS. Multivariate ANOVA or Mixed-Factor ANOVA with group, sex, and time as factors were used to test for differences, and Tukey HSD post hoc tests were used to assess the significance of differences between groups. The data are presented as mean ± standard error. For all tests, p < 0.05 was considered significant.

## Results

To investigate whether there are differences in memory facilitation in male and female rats when they are presented with two separate fear conditioning events, we subjected rats to an initial auditory fear conditioning session followed by a visual conditioning session 1 hour or 24 hours later. 48 h after the final training session, fear to the light and tone cue was tested on consecutive days **(Fig 1A).** First, we compared freezing levels in rats during the auditory fear conditioning session by taking an average of the freezing levels during each of the CS periods **Fig. 1(B, D).** A mixed factor ANOVA with trial as a within-subject factor and group, time, and sex as between-subject factors was performed. Results from this test showed a significant effect of trial (F (1,203) = 252.47, p <.01) indicating that freezing behavior increased across the two CS trials. There was also a trial by group (F (2,203) = 7.90, p <.01) and a trial by sex interaction (F (1,203) = 252.470, p <.05), presumably driven by lower levels of freezing behavior during the second trial in the shock alone group and in female rats. Finally, between-subject contrasts indicate that there is a significant effect of both group (F (2,203) = 15.69, p <.01) and sex (F (1,203) = 4.14, p <.05). Tukey HSD post hoc tests confirmed that the shock-alone group is significantly different than from both the tone-shock (p < .01) and no light-shock (p < .01) groups. These data indicate, not surprisingly, that rats trained with an unsignaled shock show less freezing than auditory fear conditioned rats during the period of time in which the discrete CS occurred.

**Figure 1.**
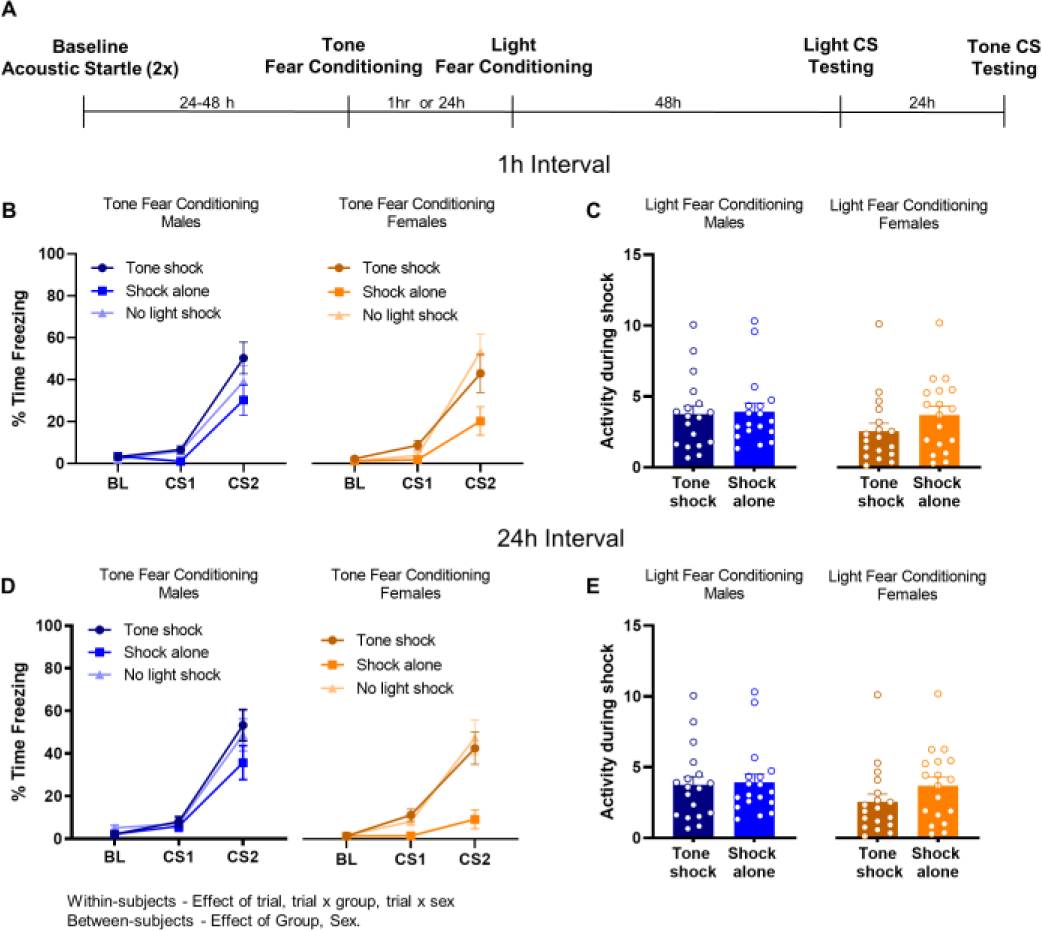
(A) Timeline of experiment. Female (N = 108, N=18 for each group) and male (N =107, N=17-19 per group) rats were given an initial tone fear conditioning session consisting of either two signaled, or two unsignaled shocks followed either 1 hour or 24 hours later by a single light-shock pairing or context exposure. Long-term memory to the tone and light cue was tested 48 h later, with the light cue tested first and the tone cue 24 h later. Freezing behavior for female and male rats during the auditory fear conditioning trial, showing average freezing behavior during the baseline (BL) period and during the two conditioning trials. Rats trained with the 1 hour (B) and 24 hour interval (D) are depicted separately for ease of comparison. Shock reactivity levels for female and male rats during the visual fear conditioning trial in rats trained with the 1 hour (C) and 24 hour (E) interval. Text below summarizes the significant effects. Error bars = +/-SEM; Dots = individual means.

One or twenty-four hours after auditory fear conditioning, rats in the tone-shock and shock-alone groups received a single light-shock pairing during visual fear conditioning, while rats in the no light-shock group were exposed to the same context but in the absence of light-shock pairing **Fig. 1(C, G)**. Although we do not obtain a measure of fear during the visual fear conditioning session, we did test whether reactivity to shock differed across conditions. A three-way ANOVA revealed no significant effect of group, time, or sex. Likewise, there were no significant interaction effects.

Two days after the auditory and visual fear conditioning sessions, all rats were tested for fear to the light cue **Fig. 2 (A, C)**. We used a three by two ANOVA to analyze the effects of sex, time, and group on fear-potentiated startle responses during the test session. Results from this analysis showed that there was a significant effect of group (F (2,203) = 4.65, p <.05) and sex (F (1,203) = 4.37, p <.05). Post hoc tests showed that the no light-shock and tone-shock groups were significantly different (p < .05). The comparison between no light-shock and shock-alone did not reach significance (p = .08). Finally, there was also a time by sex interaction (F (1,203) = 4.65, p <.05), but no other interactions between factors, indicating that females show less fear to the light cue when the two training events are separated by a day.

**Figure 2.**
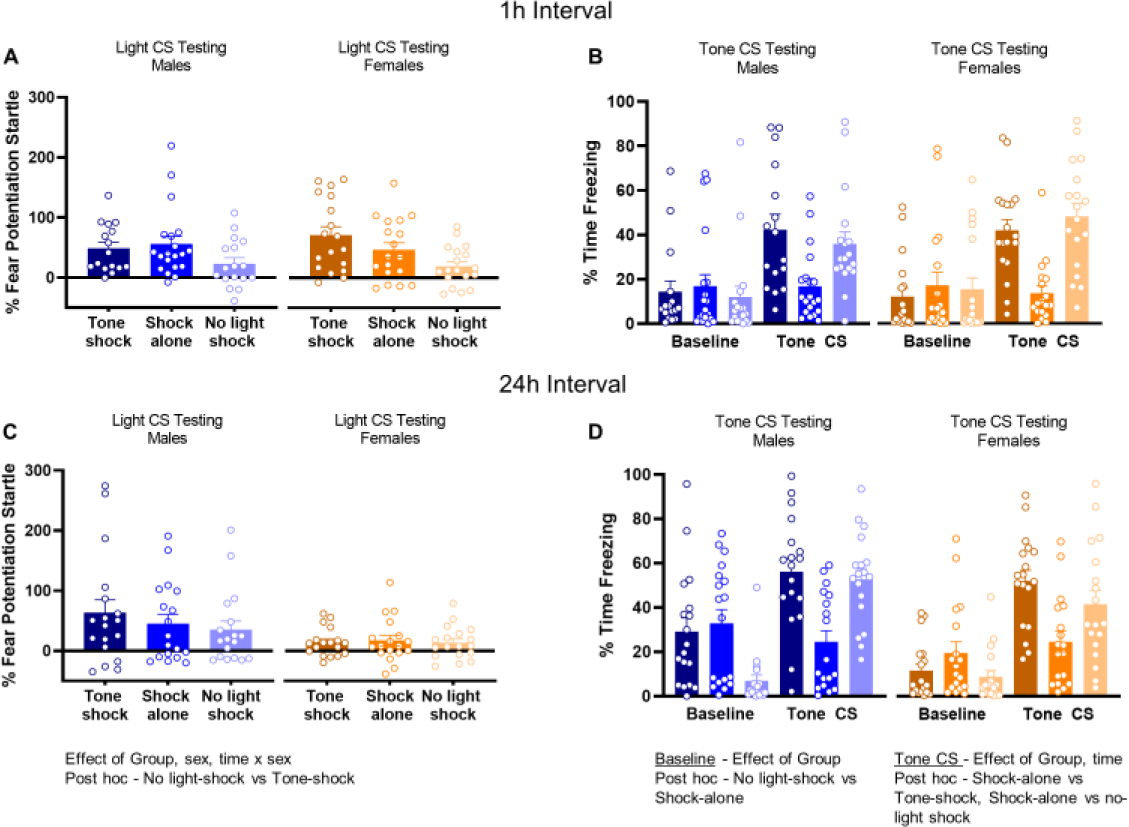
Fear behavior during the testing to the light (A, C) and auditory cues (B, D). Long-term memory to the light cue was tested first, and the tone cue 24 hours later. Fear potentiated startle levels to the light cue during the test session in female and male rats trained with the 1 hour interval (A) or 24 hour interval (C). Freezing levels during the baseline period and to the auditory tone in the same groups. Text below summarizes the significant effects.

Rats were tested 24 h later to the tone cue **Fig. 2(B, D)** in a novel context. We first compared average freezing levels across conditions during the baseline period 6 minutes before the tone was presented, which is a test of fear to the novel context in which tone testing occurred. There was a significant difference between groups (F (2,203) = 5.08, p <.01), with post hoc tests showing a significant difference between the shock-alone and no-light shock groups (p <.01). That the no-light shock group showed lower levels of freezing during the baseline period is expected given that they had only been trained once prior, while the other groups had been trained twice in difference contexts. Next, we compared average freezing levels to the tone using a MANOVA. Results from this analysis uncovered a significant effect of group (F (2,203) = 35.17, p <.01) and time (F (1,203) = 8.28, p <.01), with the effect of time reflecting higher freezing levels in rats trained with the 24 hour interval. Post hoc tests revealed that freezing during the tone in the shock-alone group, which received unsignaled shocks during the initial training session, was significantly different that both the tone-shock (p <.01) and no light-shock (p <.01) groups.

Finally, we also performed additional analyses on the light CS testing data to address two possible features of the data which might have affected the fear potentiated results depicted in **Figure 2 (A, C)**. First, we assessed the level of fear prior to presentations of the CS (i.e., baseline fear) during testing for the light cue. We did so by computing the percent change in startle responses during the startle-alone trials prior to light presentation during the CS test from the baseline startle values obtained prior to the fear conditioning sessions. A MANOVA with group, sex, and time as factors revealed no main effects and no interactions on the baseline fear measure (**Fig 3. A, C**). Coupled with the observed overall low levels of baseline fear, these results suggest that responses during baseline was unlikely to have affected fear to the light CS. Second, given that the percent change value used to measure fear potentiated startle is computed by expressing startle during the CS as a percentage of startle during the startle-alone trials presented between CS trials, differences in startle-alone values across conditions might have affected the fear potentiated startle values. To test this, we computed ‘ITI Fear’ by calculating a percent change score in startle responding during the startle alone trials after CS presentation from the alone trials prior to CS presentation. Results from a MANOVA on these data showed a significant effect of sex, with females showing higher levels of ITI fear **Figure 3 (B, D)**, but no other main effects and no significant interaction between factors. The absence of a time by sex interaction in ITI fear suggests that differences in ITI fear are unlikely to account for the finding that females exhibit little evidence of facilitation when two fear conditioning events are separated by 24 hours.

**Figure 3.**
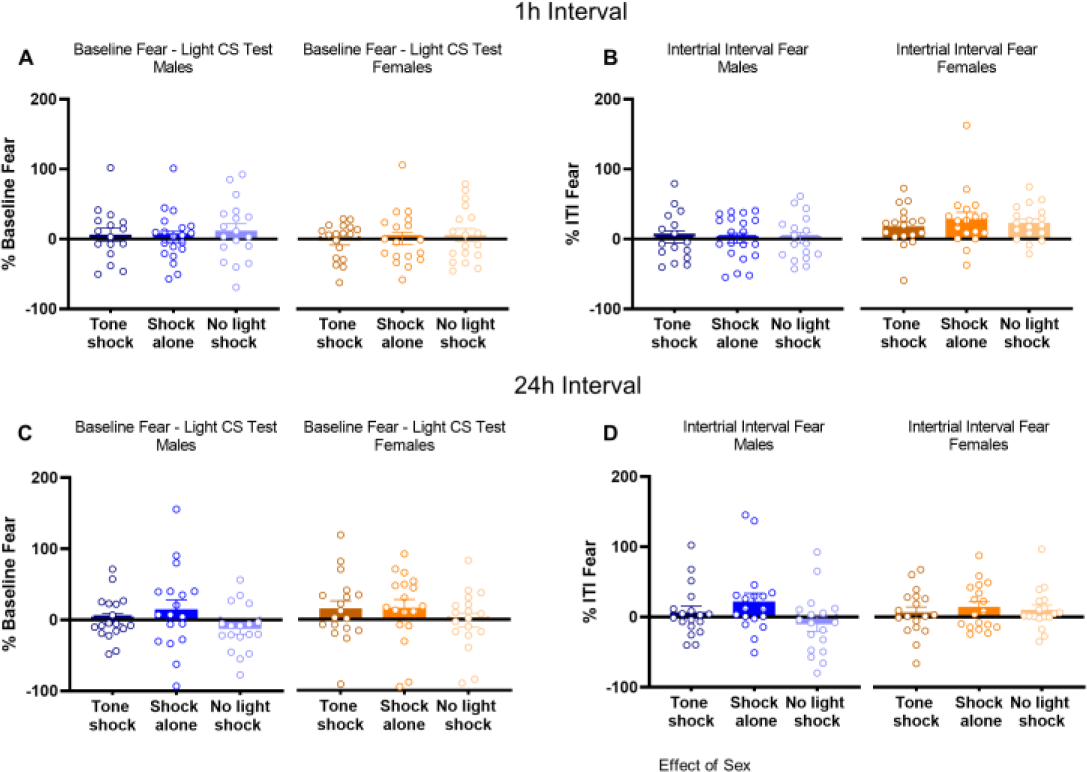
Baseline fear during the light CS test in males and females trained with a 1 hour (A) or 24 hour (C) interval. Fear during the intertrial interval (ITI) in the light CS test session in rats trained with a 1 hour (B) or 24 hour (D) interval.

## Discussion

The present results show a sex difference in the ability of auditory fear conditioning to facilitate subsequent visual fear learning. This difference was dependent on the interval between training sessions such that females displayed significantly less facilitation of visual fear conditioning when the interval was 24 hours, while both males and females showed equivalent and robust levels of facilitation when two training sessions were separated by 1 hour. Our findings also show that contextual fear conditioning did not have the same facilitatory effect, as the level of fear in both sexes trained with unsignaled shocks did not differ from rats that did not receive visual fear conditioning, which is consistent with prior work (Parsons and Davis, 2012; Cole and Parsons, 2019). However, the trend towards a difference in the rats trained with contextual fear conditioning in the current study leaves open the possibility that the right amount or intensity of contextual fear conditioning might facilitate later learning. Altogether, these findings demonstrate that there are fundamental differences between sexes in how prior aversive experience alters subsequent fear learning and adds important knowledge about the factors that influence the ability of prior learning to alter subsequent memory formation.

Our current results showed that prior auditory fear conditioning is capable of facilitating subsequent visual fear learning when rats are later presented with a single pairing of light and shock. This might appear to be inconsistent with prior published work (Parsons and Davis, 2012), which showed that facilitation only occurred if the first and second training sessions were identical and not if the shock was presented alone or paired with an auditory cue during the first session. However, several factors likely account for this difference, including the fact that during the initial conditioning episode, the two studies differed with respect to the number and intensity of shocks, with the current work using more trials and a higher intensity shock, both of which would likely favor the development of subsequent facilitation. Additionally, in the prior work, both training sessions occurred in the same context, whereas for the study presented here, the two conditioning sessions took place in distinct contexts. In fact, more recent work from our lab (Lee et al., 2018) showed that auditory fear conditioning is capable of facilitating subsequent visual fear learning when multiple trials of auditory fear conditioning are administered in a unique context. Thus, it is likely that some combination of the difference in context and the intensity of initial training between the two studies accounts for discrepant results.

Importantly, two aspects of our data suggest that reduced fear to the light cue in females trained with a 24 hour interval between conditioning sessions is not simply the result of an overall reduction in the expression of fear in females. First, females trained with a 1 hour interval between sessions showed no difference in fear to the light cue as compared to males. Prior work comparing males and females on fear potentiated startle using more typical training procedures is somewhat equivocal, although some reports have found differences (de Jongh et al., 2005), others have reported no differences between males and females (Voulo and Parsons, 2017; Zhao et al., 2018; Russo and Parsons, 2021). Second, there were also no differences between sexes in freezing when rats were tested to the auditory cue, consistent with a number of previous findings (Maren et al., 1994; Milad et al., 2009a; Barker and Galea, 2011; Gruene et al., 2015; Fenton et al., 2016; Voulo and Parsons, 2017), and again suggesting that our results do not simply reflect a difference in the expression of fear between males and females.

An ongoing goal of the work in our lab has been to determine the neural mechanisms that allow prior fear conditioning to alter the strength of subsequent fear learning. At least in males, there is good evidence that the facilitation of subsequent learning is dependent on some of the same molecular mechanisms as memory consolidation for the initial training. For example, our prior work has identified both the protein kinase A and mitogen-activated protein kinase signaling in the basolateral amygdala as being necessary for initial learning to facilitate subsequent fear conditioning (Parsons and Davis, 2012; Parsons, Walker, and Davis, 2016; Cole et al., 2019). The cyclic-AMP response element binding protein has also been implicated by our prior work (Parsons and Davis, 2012) and others studying the neural basis of fear memory linking (Rashid et al., 2016). It is unknown whether the same signaling mechanisms are involved in shorter-term facilitation effects in females, although it would not be surprising if they do, given the evidence that the molecular (Datchler et al., 2011; Devulapalli et al., 2021; Farrell et al., 2021; Martin et al., 2021) and neural circuit mechanisms (Gruene et al., 2015; Keiser et al., 2017; Urien and Bauer, 2022) underlying typical fear learning and the expression of fear differ in males and females (for review, Tronson and Keiser, 2019). Another consideration to take into account is that in many of the prior studies examining the molecular mechanisms of memory facilitation, the time course of the behavioral facilitation effects are congruous with the molecular changes, such that facilitation occurs when the second training event takes place within the time window of molecular consolidation engaged by the initial experience (reviewed in Parsons, 2018). However, as of yet, it is not clear which neural mechanisms might allow for the relatively long time course by which prior auditory fear learning is able to facilitate subsequent visual fear conditioning. Based on the findings here, it would be expected that the mechanisms that bridge this longer time course would be differentially engaged in males and females.

The findings reported here add to our understanding of the factors supporting the ability of prior fear conditioning to facilitate subsequent learning. Together with our prior work, we now know that both identical and non-identical conditioning events can facilitate subsequent fear learning, that there is an optimal time window for this facilitation effect to occur (Parsons and Davis, 2012; Lee et al., 2018), and that although males show facilitation when the two events are spaced at longer intervals, auditory fear conditioning does not facilitate subsequent visual fear conditioning in females if the two training sessions are separated by 24 hours. One possibility is that facilitation at the longer intervals is supported by hippocampus-dependent contextual processes, which would be consistent with other data showing that male rodents exhibit higher levels of contextual fear than females (Maren et al., 1994; Chang et al., 2009; Barker and Galea, 2011) and generally consistent with a larger literature on sex and hippocampal plasticity (McEwen and Milner, 2007; Koss and Frick, 2017). However, the fact that many studies have found no sex difference in contextual fear (Kosten et al., 2006; Dachtler et al., 2011; Keiser et al., 2017), and our recent findings showing the deficit is specific to freezing behavior (Russo and Parsons, 2021), suggest that there are unknown factors that dictate whether a sex difference is observed. Additionally, in the present study, we showed that when rats are trained without a discrete CS, the ability of initial learning to facilitate subsequent fear conditioning is lost. Although the trend towards a difference makes this finding somewhat ambiguous, the lack of an effect with context fear conditioning would seem to suggest context might not govern facilitation unless it is the case that the contextual processing differs based on whether or not shock is preceded by a discrete CS, for which there is some evidence (Phillips and LeDoux, 1994). Nonetheless, it would be of interest to systematically test the role of context in our paradigm.

We also found that females showed overall higher levels of fear during the intertrial interval during testing to the light cue. Because fear to the light was computed by expressing startle amplitudes during light-startle trials as a percentage of the startle-alone trials presented during the ITI, differences between groups in ITI fear might confound the measure of fear to the light CS. Thus, the fact that females showed higher levels of ITI fear might have artificially reduced the computed levels of fear to the CS in females. However, differences in fear during the ITI cannot explain the finding that females trained with the two conditioning events separated by 24 hours show little evidence of facilitation, as higher levels of ITI fear in females were not specific to the 24 hour interval (i.e., there was no interaction between sex and training interval). Although behavior during the ITI at testing is not often reported in most studies of fear learning, some recent findings have reported females exhibiting higher levels of fear during the ITI at later points during training (Borkar et al., 2020). Although it is been widely reported that the ITI is a critical variable affecting conditioning, there are relatively few data pertaining to what fear during the ITI at testing reflects. Some possibilities might include residual fear from the CS, fear in anticipation of an upcoming trial (Zielinski and Nikolaev, 1997), or context fear as a result of a CS-context association made during training (Quinn et al., 2002). More work is needed to understand the nature of ITI fear and whether or not consistent sex differences are observed during the ITI.

The results reported have some noteworthy limitations, including the fact that we did not track estrous in the females. The extant data are mixed with an effect of the estrous cycle often being dependent on which phase of fear conditioning is under study (i.e. training, testing, or extinction), whether the study used a cued or contextual fear paradigm, how fear behavior was measured, and whether or not rodents were trained and tested in the same phase (Milad et al., 2009; Lynch et al., 2013, Chang et al., 2009; Gresack et al., 2009; Fenton et al., 2016; Zhao et al., 2018, Voulo and Parsons, 2019; Blair et al., 2022). Future studies are needed to determine if estrous cycle phase affects the facilitation of learning by prior fear conditioning. Another limitation is related to the possibility that parametric differences between the two different training experiences affected our findings. The two training events had different number of trials, CS modalities, UCS intensity, and response measurement. Most of these differences are out of necessity, as we have described before (Voulo and Parsons, 2019), and also to create conditions that favor subthreshold learning, which is necessary for being able to detect the facilitation of learning. For these parametric differences to have affected our main findings, they would have had to have effects that are specific to females trained with a 24 hour interval.

In conclusion, our results show that there are fundamental differences between males and females in how prior aversive experience alters subsequent fear learning. Male rats trained with two cued fear conditioning events separated by 60 minutes or 24 hours showed evidence that the first conditioning event was able to facilitate learning of the second. While female rats showed facilitation when the training events were separated by 60 minutes, there was no evidence of facilitation in females trained with the two events separated by 24 hours. In addition, we found that contextual fear conditioning was unable to facilitate subsequent fear learning at the two intervals tested and in either sex. Our findings provide important knowledge about the factors that support the ability of prior fear conditioning to facilitate subsequent fear learning, and they also provide the foundation for study into the neurobiological mechanisms of sex differences in fear memory facilitation.

## Funding Statement

This research was supported by startup funds from Stony Brook University, The Stony Brook Foundation, and grants R21 MH121772 (to R.G.P) from the U.S. National Institutes of Health.

## Notes

### Competing Interest Statement

The authors have declared no competing interest.

## References

Blair, R.S., Acca, G.M., Tsao, B., Stevens, N., Maren, S., and Nagaya, N. (2022). Estrous cycle contributes to state-dependent contextual fear in female rats. Psychoneuroendocrinology 141, 105776.

Barker, J.M., and Galea, L.A. (2010). Males show stronger contextual fear conditioning than females after context pre-exposure. Physiol Behav 99, 82–90.

Borkar, C.D., Dorofeikova, M., Le, Q.E., Vutukuri, R., Vo, C., Hereford, D., Resendez, A., Basavanhalli, S., Sifnugel, N., and Fadok, J.P. (2020). Sex differences in behavioral responses during a conditioned flight paradigm. Behav Brain Res 389, 112623.

Cole, K.E., Lee, J., Davis, M., and Parsons, R.G. (2019). Subthreshold Fear Conditioning Produces a Rapidly Developing Neural Mechanism that Primes Subsequent Learning. eNeuro 6.

Chang, Y.J., Yang, C.H., Liang, Y.C., Yeh, C.M., Huang, C.C., and Hsu, K.S. (2009). Estrogen modulates sexually dimorphic contextual fear extinction in rats through estrogen receptor beta. Hippocampus 19, 1142–1150.

Dachtler, J., Fox, K.D., and Good, M.A. (2011). Gender specific requirement of GluR1 receptors in contextual conditioning but not spatial learning. Neurobiol Learn Mem 96, 461–467.

de Jongh RL, Geyer MA, Olivier B, Groenink L. 2005. The effects of sex and neonatal maternal separation on fear-potentiated and light-enhanced startle. Behav Brain Res 161, 190–196.

Devulapalli, R., Jones, N., Farrell, K., Musaus, M., Kugler, H., Mcfadden, T., Orsi, S.A., Martin, K., Nelsen, J., Navabpour, S., O’donnell, M., Mccoig, E., and Jarome, T.J. (2021). Males and females differ in the regulation and engagement of, but not requirement for, protein degradation in the amygdala during fear memory formation. Neurobiol Learn Mem 180, 107404.

Farrell, K., Musaus, M., Navabpour, S., Martin, K., Ray, W.K., Helm, R.F., and Jarome, T.J. (2021). Proteomic Analysis Reveals Sex-Specific Protein Degradation Targets in the Amygdala During Fear Memory Formation. Front Mol Neurosci 14, 716284.

Fenton, G.E., Halliday, D.M., Mason, R., Bredy, T.W., and Stevenson, C.W. (2016). Sex differences in learned fear expression and extinction involve altered gamma oscillations in medial prefrontal cortex. Neurobiol Learn Mem 135, 66–72.

Gresack, J.E., Schafe, G.E., Orr, P.T., and Frick, K.M. (2009). Sex differences in contextual fear conditioning are associated with differential ventral hippocampal extracellular signal-regulated kinase activation. Neuroscience 159, 451–467.

Gruene, T. M., Roberts, E., Thomas, V., Ronzio, A., & Shansky, R. M. (2015). Sex-specific neuroanatomical correlates of fear expression in prefrontal-amygdala circuits. Biol Psychiatry, 78, 186–193.

Keiser, A.A., Turnbull, L.M., Darian, M.A., Feldman, D.E., Song, I., and Tronson, N.C. (2017). Sex Differences in Context Fear Generalization and Recruitment of Hippocampus and Amygdala during Retrieval. Neuropsychopharm 42, 397–407.

Kessler RC, Sonnega A, Bromet E, Hughes M, Nelson CB. 1995. Posttraumatic stress disorder in the national comorbidity survey. Arch Gen Psychiatry 52, 1048–1060.

Kilpatrick DG, Resnick HS, Milanak ME, Miller MW, Keyes KM, Friedman MJ. 2013. National estimates of exposure to traumatic events and PTSD prevalence using DSM-IV and DSM-5 criteria. J Trauma Stress 26, 537–547.

Koss, W.A., and Frick, K.M. (2017). Sex differences in hippocampal function. J Neurosci Res 95, 539–562.

Kosten, T.A., Lee, H.J., and Kim, J.J. (2006). Early life stress impairs fear conditioning in adult male and female rats. Brain Res 1087, 142–150.

Lee, J., Russo, A.S., and Parsons, R.G. (2018). Facilitation of fear learning by prior and subsequent fear conditioning. Behav Brain Res 347, 61–68.

Lynch, J., 3rd, Cullen, P.K., Jasnow, A.M., and Riccio, D.C. (2013). Sex differences in the generalization of fear as a function of retention intervals. Learn Mem 20, 628-632.

Maren, S., De Oca, B., and Fanselow, M.S. (1994). Sex differences in hippocampal long-term potentiation (LTP) and Pavlovian fear conditioning in rats: positive correlation between LTP and contextual learning. Brain Res. 661, 25–34.

Martin, K., Musaus, M., Navabpour, S., Gustin, A., Ray, W.K., Helm, R.F., and Jarome, T.J. (2021). Females, but not males, require protein degradation in the hippocampus for contextual fear memory formation. Learn Mem 28, 248–253.

Mcewen, B.S., and Milner, T.A. (2007). Hippocampal formation: shedding light on the influence of sex and stress on the brain. Brain Res Rev 55, 343–355.

Milad, M.R., Igoe, S.A., Lebron-Milad, K., and Novales, J.E. (2009). Estrous cycle phase and gonadal hormones influence conditioned fear extinction. Neuroscience 164, 887–895.

Parsons, R. G., & Davis, M. (2012). A metaplasticity-like mechanism supports the selection of fear memories: Role of protein kinase a in the amygdala. J Neurosci, 32, 7843–7851.

Parsons, R.G., and Ressler, K.J. (2013). Implications of memory modulation for post-traumatic stress and fear disorders. Nat Neurosci 16, 146–153.

Parsons, R.G., Walker, D.L., and Davis, M. (2016). Mechanisms underlying long-term fear memory formation from a metaplastic neuronal state. Neurobiol Learn Mem 136, 47–53.

Parsons, R.G. (2018). Behavioral and neural mechanisms by which prior experience impacts subsequent learning. Neurobiol Learn Mem 154, 22–29.

Phillips, R.G., and Ledoux, J.E. (1994). Lesions of the dorsal hippocampal formation interfere with background but not foreground contextual fear conditioning. Learning & memory 1, 34–44.

Quinn, J.J., Oommen, S.S., Morrison, G.E., and Fanselow, M.S. (2002). Post-training excitotoxic lesions of the dorsal hippocampus attenuate forward trace, backward trace, and delay fear conditioning in a temporally specific manner. Hippocampus 12, 495–504.

Rashid, A.J., Yan, C., Mercaldo, V., Hsiang, H.L., Park, S., Cole, C.J., De Cristofaro, A., Yu, J., Ramakrishnan, C., Lee, S.Y., Deisseroth, K., Frankland, P.W., and Josselyn, S.A. (2016). Competition between engrams influences fear memory formation and recall. Science 353, 383–387.

Russo, A.S., and Parsons, R.G. (2021). Behavioral Expression of Contextual Fear in Male and Female Rats. Front Behav Neurosci 15, 671017.

Tronson, N.C., and Keiser, A.A. (2019). A Dynamic Memory Systems Framework for Sex Differences in Fear Memory. Trends Neurosci 42, 680–692.

Urien, L., and Bauer, E.P. (2022). Sex Differences in BNST and Amygdala Activation by Contextual, Cued, and Unpredictable Threats. eNeuro 9.

Voulo, M. E., & Parsons, R. G. (2017). Response-specific sex difference in the retention of fear extinction. Learn Mem, 24, 245–251.

Voulo, M.E., and Parsons, R.G. (2019). Gonadal hormone fluctuations do not affect the expression or extinction of fear-potentiated startle in female rats. Behav Neurosci 133, 517–526.

Walker, D.L., and Davis, M. (1997). Anxiogenic effects of high illumination levels assessed with the acoustic startle response in rats. Biol Psychiatry 42, 461–471.

Walker, D.L., and Davis, M. (2002). Light-enhanced startle: further pharmacological and behavioral characterization. Psychopharm (Berl) 159, 304–310.

Zhao, Y., Bijlsma, E. Y., Verdouw, M. P., & Groenink, L. (2018). No effect of sex and estrous cycle on the fear-potentiated startle response in rats. Behav Brain Res 351, 24–33.

Zielinski, K., and Nikolaev, E. (1997). Changes of intertrial response rate with elapse of time after two-way avoidance trial in rats. Acta Neurobiol Exp (Wars) 57, 41–47.

